# Carbon stocks of coastal seagrass in Southeast Asia may be far lower than anticipated when accounting for black carbon

**DOI:** 10.1101/493262

**Authors:** John B. Gallagher, Chuan Chee Hoe, Yap Tzuen-Kiat, W Farahain

## Abstract

Valuing sedimentary ‘blue carbon’ stocks of seagrass meadows requires exclusion of allochthonous recalcitrant forms of carbon, such as black carbon (BC). Regression models constructed across a Southeast Asian tropical estuary predicted that carbon stocks within the sandy meadows of coastal embayments would support a modest but not insignificant amount of BC. We tested the prediction across three coastal meadows of the same region: one patchy meadow located close to a major urban centre and two continuous meadows contained in separate open embayments of a rural marine park; all differed in fetch and species. The BC/total organic carbon (TOC) fractions in the urban and rural meadows with small canopies were more than double the predicted amounts, 28 ± 1.6% and 36 ± 1.5% (±95% confidence intervals), respectively. The fraction in the rural large-canopy meadow remained comparable to the other two meadows, 26 ± 4.9% (±95% confidence intervals) but was half the amount predicted, likely due to confounding of the model. The relatively high BC/TOC fractions were explained by variability across sites of BC atmospheric supply, an increase in loss of seagrass litter close to the exposed edges of meadows, and sediment resuspension across the dispersed patchy meadow.

## Introduction

The effect of anthropogenic emissions of CO_2_ on the climate highlights the importance of quantifying and managing existing sedimentary organic carbon stocks. Lately, focus has turned to ‘blue carbon’ reservoirs – seagrass, saltmarsh, mangrove and macro-algae ecosystems (1, 2). Together, these store half the ocean’s total organic carbon (TOC), despite occupying less than 2% of the area (3). This disproportionate contribution results from their high rates of primary production and ability to enhance and stabilise accumulation of deposited litter and detritus and prevent it from remineralising, ostensibly back to CO_2_ (4, 5). Of the four, seagrass ecosystems are best placed to augment organic carbon stocks. Seagrass canopy can trap soils eroded from adjacent landscapes, which can account for as much as 50% of total sedimentary organic stocks (6) that would otherwise be remineralised across the continental shelf (7).

Recently, the traditional mass balance approach to calculating blue carbon storage has been challenged (8). It is argued that recalcitrant organic carbon produced outside an ecosystem does not require protection from remineralisation. Consequently, such deposits cannot be counted as a service in the mitigation of greenhouse gas emissions. Black carbon (BC) is an example of an ‘allochthonous recalcitrant’; it is formed during the incomplete combustion of biomass and fossil fuels, for which Southeast Asia is a global hotspot (9). However, the BC content of coastal seagrass sediments is unknown and thus too is the extent of bias in stock estimates. Nonetheless, predictions of ∼ 18 ± 3% (±95% confidence interval) have been made using a Southeast Asian (Sabah, Malaysia) estuarine TOC-BC regression model (8). The model’s parameters (i.e. large BC intercept) indicate that high BC/TOC fractions are likely in relatively low organic content sandy sediments of coastal meadows in regions where atmospheric BC is pervasive over that of soil erosion (i.e. a small regression slope). The slope is determined by the accumulated BC within the soil profile and is tempered by the edaphic organic content of the soil itself (8). Taken together, such regressions give first-order predictive estimates of meadow BC/TOC.

The accuracy of estuarine model’s BC/TOC coastal predictions around this region, while moderate, may be confounded by failure to take into account the relatively efficient retention of seagrass litter within estuaries. In more exposed coastal systems, most of the litter across a meadow or close to its exposed boundary can be lost to shelf waters (10–12). It is also possible that increased turbulence closer to the exposed edges or across more patchy seagrass seascapes may lead to localised sediment resuspension. This may further exacerbate the remineralisation of more labile organic fractions, thereby amplifying the importance of BC to the TOC stock.

This study examines the above estuarine predictions and likely controlling factors (i.e. litter loss and sediment resuspension) across coastal meadows in Southeast Asia (Sabah, Malaysia). We measured the variance in TOC, BC, canopy and sedimentary variables along transects across and between three subtidal (0.5–1 m) seagrass meadows, two relatively pristine and one impacted by surrounding developments.

## Materials and Methods

Transects were set behind each other and perpendicular to the boundary exposed to prevailing monsoons. Although replication of the meadows within the relatively pristine environs of the Tun Mustapha Marine Park, at Limau-Limauan (LL) and Bak-Bak (BB), was not possible, we controlled for good coverage. Confounding effects were reduced by selecting extreme examples of canopy size (13) covariant with extremes of wind fetch (14). LL supported *Enhalus acoroides* as a large climax species, and BB supported a mix of smaller canopy pioneer species, including *Cymodocea rotundata* and *Halodule pinifolia* (supplemental Figure S1). The third meadow, chosen for its sparse, patchy configuration, was located near the University of Malaysia’s Outdoor Development Centre beach (OD), which is under pressure from land development across the urban environs of Sepanggar Bay (15). The area has a relatively moderate wind fetch and canopy size. It sustains meadows comprising *Cymodocea serrulata, Halodule uninervis* and *Halophila ovalis* and isolated small stands of *E. acoroides*, which assisted in reducing confounding comparisons with the rural systems (supplemental Figure S1). Details of sediment collection, measurements of canopy and sediment parameters and TOC and BC analysis are found in the electronic supplementary information.

## Results

### Meadow parameters

The meadow canopies exhibited clear height differences, attributable to species characteristics and relative wind fetch (supplemental Figure S1), in the order of LL >> BB < OD (Table 1). Differences in canopy coverage between meadows LL and BB in the relatively pristine marine park and OD in the relatively polluted Sepanggar Bay were as expected (LL ≈ BB >> OD, Table 1). Furthermore, significant differences in both height and coverage between transect pairs was only apparent for OD, reflecting its sparse, patchy configuration.

**Table 1.**
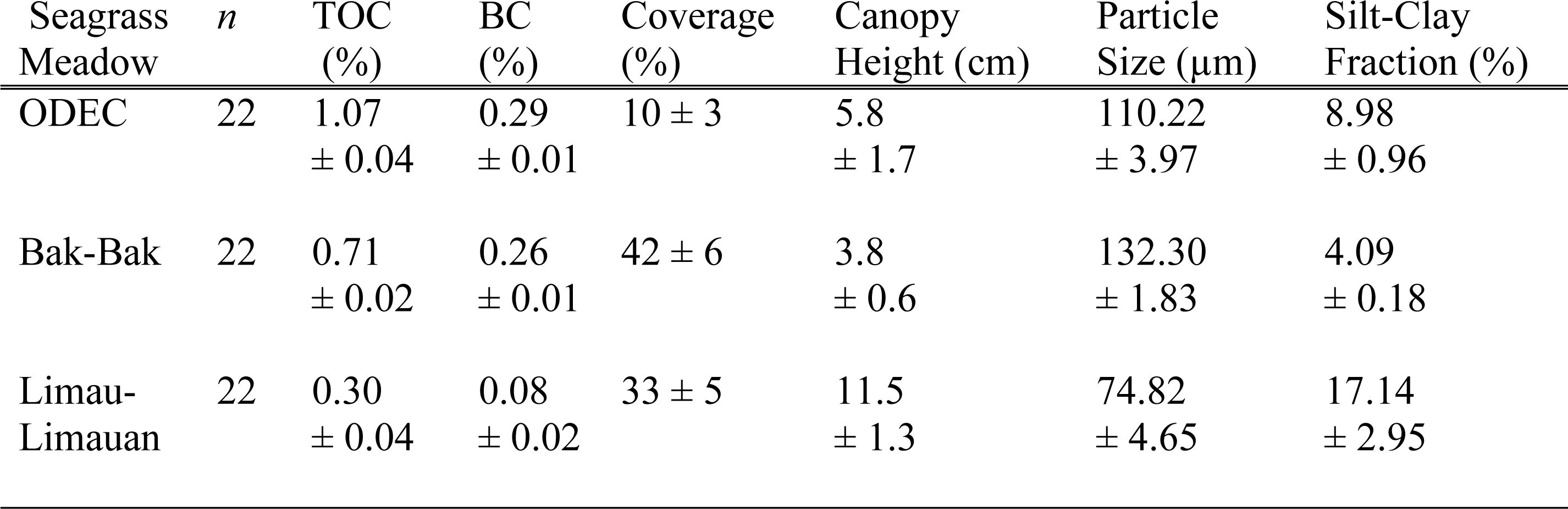
Mean and standard errors of total organic carbon (TOC) and black carbon (BC) surface sediment contents (as dry mass) and seagrass canopy and sediment parameters for three Sabahan seagrass meadows.

| Seagrass Meadow | *n* | TOC (%) | BC (%) | Coverage (%) | Canopy Height (cm) | Particle Size (μm) | Silt-Clay Fraction (%) |
| --- | --- | --- | --- | --- | --- | --- | --- |
| ODEC | 22 | 1.07<br>± 0.04 | 0.29<br>± 0.01 | 10 ± 3 | 5.8<br>± 1.7 | 110.22<br>± 3.97 | 8.98<br>± 0.96 |
| Bak-Bak | 22 | 0.71<br>± 0.02 | 0.26<br>± 0.01 | 42 ± 6 | 3.8<br>± 0.6 | 132.30<br>± 1.83 | 4.09<br>± 0.18 |
| Limau-Limauan | 22 | 0.30<br>± 0.04 | 0.08<br>± 0.02 | 33 ± 5 | 11.5<br>± 1.3 | 74.82<br>± 4.65 | 17.14<br>± 2.95 |

The most striking difference in sediment characteristics was between LL, where the longer *Enhalus* canopy species supported higher silt-clay fractions and a significantly smaller mean particle size, and BB and OD, which were sandier (Table 1). However, these dissimilarities do not reflect inter-meadow variability between transect pairs, which showed significant differences only for OD (Table 2).

**Table 2:**
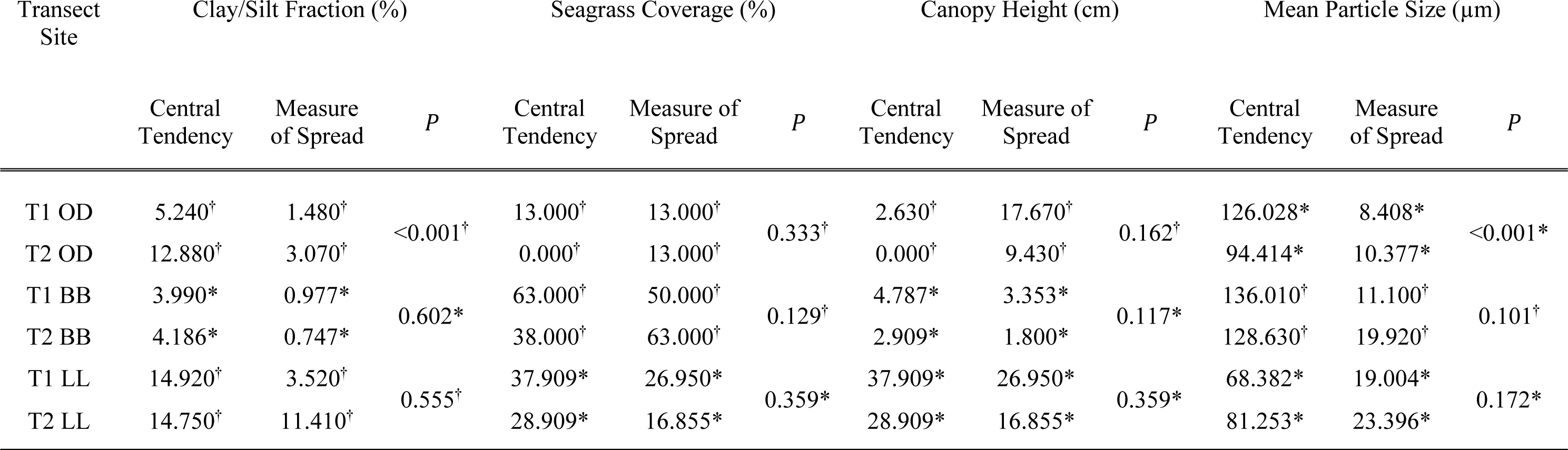
Clay/silt fraction, seagrass coverage, canopy height and mean particle size of each studied transect. Transects labelled T1 correspond to the inner transect, whereas T2 corresponds to the outer transect. †: Calculated using a Mann-Whitney rank sum test as the data were not normally distributed according to the Shapiro-Wilk normality test (*P* < 0.050); central tendency and measure of spread are the median and the interquartile range. *: Calculated using a t-test as the data were normally distributed according to the Shapiro-Wilk normality test (*P* ≥ 0.050); central tendency and measure of spread are mean and standard deviation.

Differences in both TOC and BC content between meadows (LL << OD > BB) were inverted compared to expectations regarding percentage of slit-clay content (i.e. LL >> OD < BB; Table 1). Interestingly, it was noted that the mean particle sizes within OD’s patchy stands were much greater than within adjacent bare patches of the meadow: 8.1±2.4 µm and 107.5±19.0 µm (±95% CI), respectively, but with no statistical significance between their TOC medians (*P =* 0.53; see supplemental Table S1). For LL and BB but not OD, the outer transect showed, on average, lower values for both TOC and BC, a difference which became greater and increasingly significant across LL (Figure 2a, Table 2).

### Black carbon to TOC variability

Overall, BC represented a significantly larger fraction of TOC in all three meadows than in the upper silty-mud and lower silty-sand sediments of the Salut–Mengkabong estuary (Figure 1). For the urban and rural small-canopy meadows (OD and BB), BC was 28±1.6% and 36±1.5% (±95% confidence intervals) of TOC, respectively (Table 1). The BC/TOC values of the large-canopy meadow (LL) were comparable, 26±4.9% (±95% CI; Table 1), despite large variability. This variability appears to originate with two high TOC and BC outliers (Figure 2b) located at the start of each transect (see supplemental Table S1). Furthermore, the larger BC/TOC fractions between the estuary and coastal regions were also reflected in their adjacent bare sediments.

**Figure 1.**
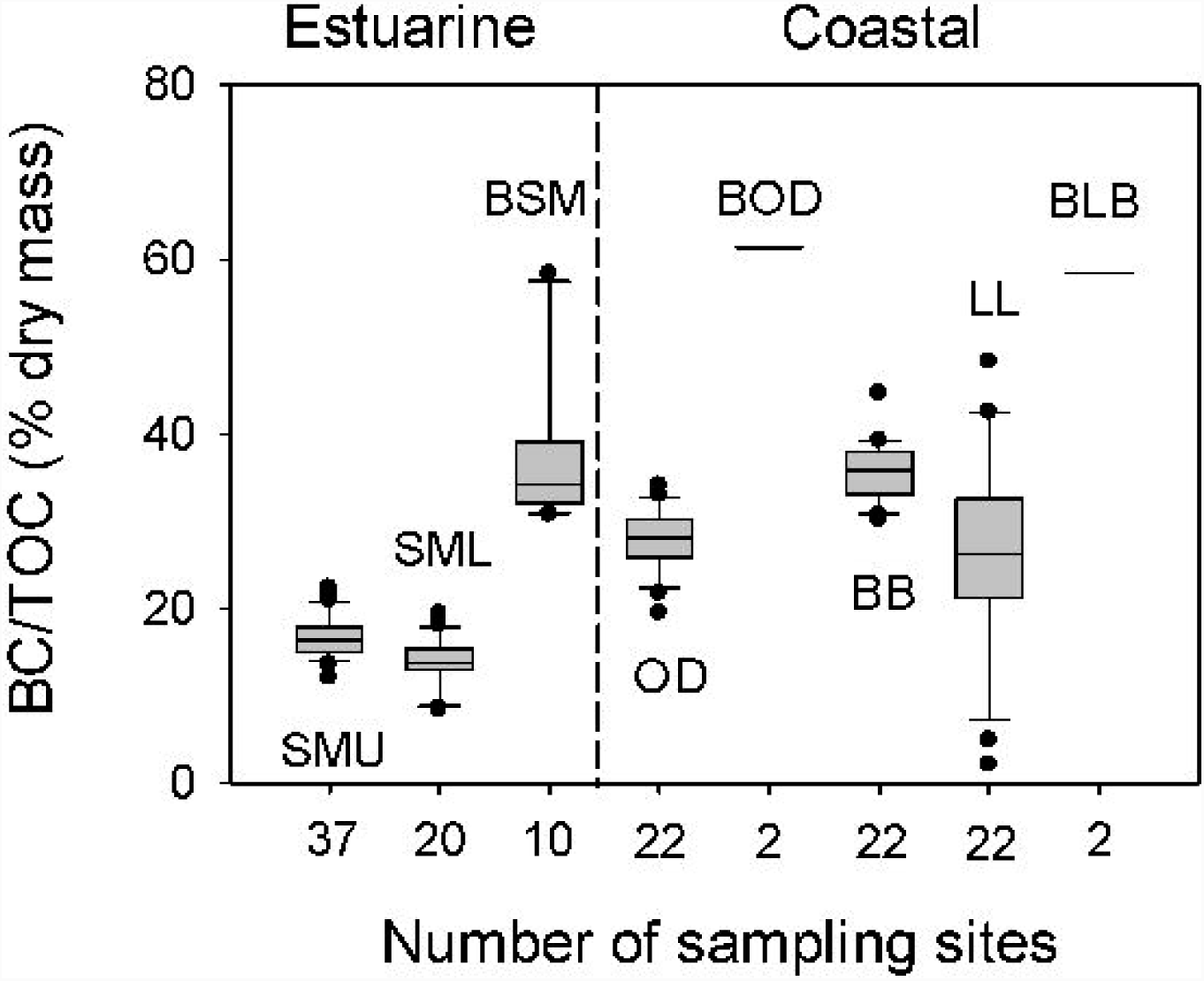
BC/TOC fractions for seagrass sediments of upper (SMU) and lower (SML) Salut– Mengkabong Estuary and the coastal seagrass meadows at ODEC beach (OD), Bak-Bak (BB) and Limau-Limauan (LL). BSM, BOD and BLB indicate the fraction from bare sediments outside the meadows within the estuary, at OD, and at LL and BB combined, respectively. The box plot shows the median, 25% and 75% quartiles, 95% confidence limits and outliers. Data for SMU and SML were compiled from supplementary material (5) in accordance with the Open Access licence http://creativecommons.org/licenses/by/4.0/.

**Figure 2.**
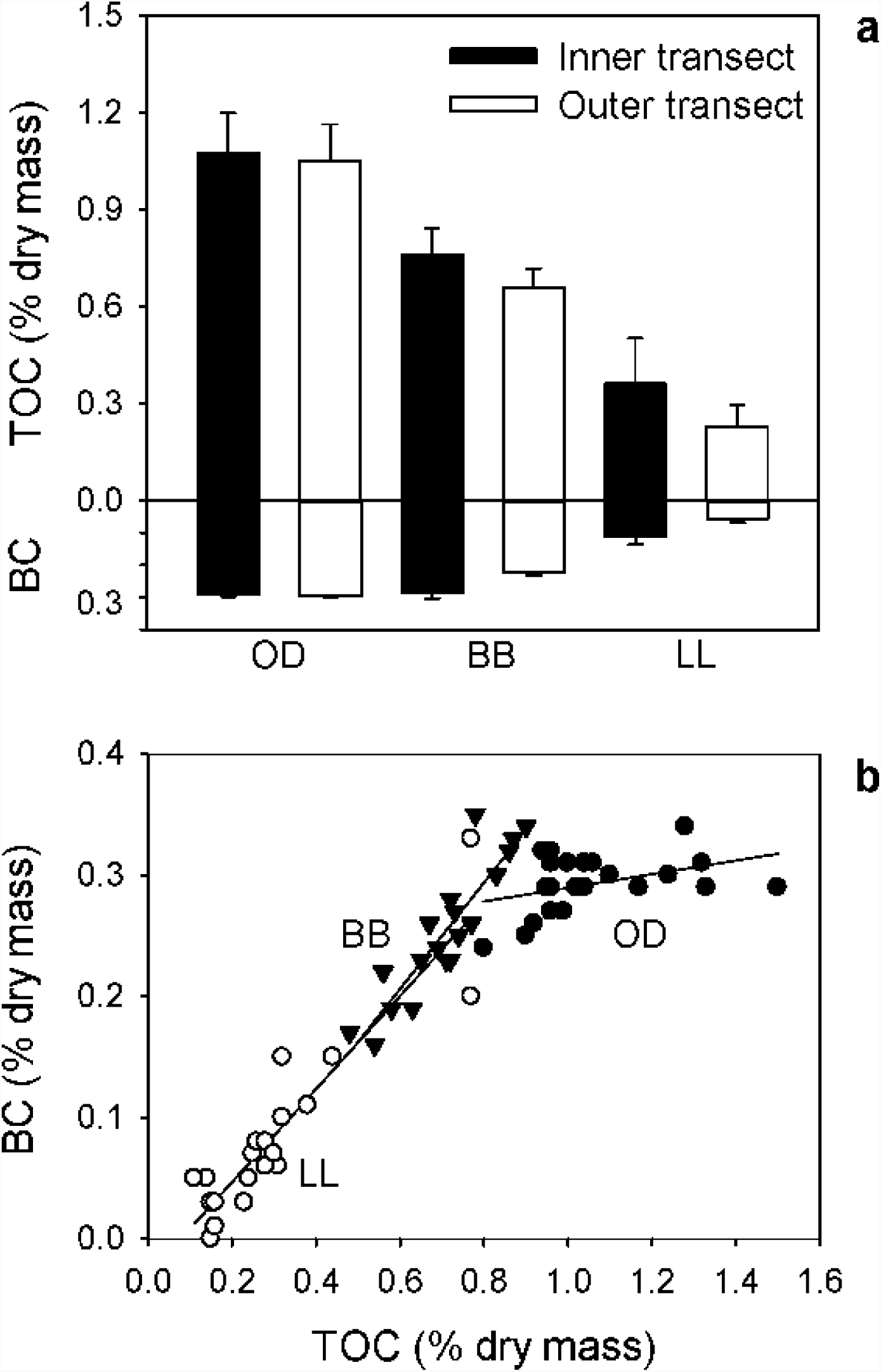
(**a**) Total organic carbon (TOC) and black carbon (BC) content of meadows at ODEC beach (OD), Bak Bak (BB) and Limau Limauan (LL) averaged across inner and outer meadow transects. Error bars represent 95% confidence limits. Significant differences between transect TOC and BC means were found for LL (*P* < 0.001, *P* < 0.001) and BB (*P* < 0.07, *P* < 0.001). A power analysis (80%) indicated that determining TOC between BB transects would require another two or three samples to resolve differences in the mean from a possible Type 1 error. (**b**) Ordinary least squares regressions for BC with TOC across the three meadows. The probability of the regression intercepts and slopes between BB and LL are significantly similar (ANCOVA *P* > 0.38) and not significantly similar to OD (ANCOVA *P* < 0.0001).

In all cases, the variability was part of a linear response. For BB and LL, the relationship between TOC and BC was identical in terms of both their proportional response and the locations of the intercept close to the origin (Figure 2b). In contrast, BC content in OD was relatively invariant with increases in TOC [*R*^*2*^ = 0.16, slope = 0.06±0.08 (±95% CI)], resulting in a BC intercept significantly greater than zero [0.23%±0.168 (±95% CI)]. Furthermore, their BC/TOC estimates were approximately half than the estuarine model could have predicted (8) from their mean TOC for the relatively small-canopy meadows, BB and OD [BB: model 17.4±2.7%; measured 36.7±0.8% (±95% CI); OD: [model 13.0±1.8%; measured 30.0±0.8% (±95% CI)], and more than double for the large-canopy meadow, LL [LL model 46.4±9.4%; measured 26.1±2.5% (±95% CI)].

## Discussion

All the coastal meadow sediments supported greater BC/TOC fractions than those found in the nearby estuary. Comparisons with the estuarine model predictions, however, were mixed. As for BB and OD, we expected the BC/TOC fractions to be larger than predicted for open embayments for a corresponding mean TOC and consistent with a greater loss of the labile fraction (see Introduction). This was not the case for the larger canopy *Enhalus sp.* of LL. This disparity may be the result of estuarine model confounding. Unlike the clear waters and sandier environs of LL, the estuarine *Enhalus sp.* meadow was found solely rooted in highly organic sediments in the upper and relatively turbid parts of the estuary (8).

The mechanism by which BC/TOC fractions have seemingly been elevated within coastal embayments can best be distinguished via comparisons of intra-meadow transects in relation to sedimentary parameters, supported by patterns consistent with the relative canopy sizes. As expected, the more exposed outer transects for both LL and BB supported lower sedimentary TOC. Considering that the transect pairs have nearly identical sedimentary parameters, falling rates of resuspension and subsequent oxidation closer to the meadows’ exposed edges is not a likely explanation. Neither do the contributions of BC from adjacent bare sediments explain the data patterns. The net deposition of these sediments is likely to be greater across the inner transects, as turbulence is increasingly attenuated (17). This should reduce rather than increase the TOC of the inner transects. Thus, a model that describes increases in BC with TOC, converging towards a positive TOC intercept close to its origin is more consistent with additional organic carbon not associated with BC such as soil washout. How much litter and soil make up these meadow’s sediments is unknown and beyond the scope of this study. The alternative explanation is reduction in the loss of deposited litter away from the exposed edge due to greater attenuation of currents by the canopy. This contention is further supported, though a weak inference, by a larger difference and increasing significance in the TOC variance for the larger canopy species of LL.

In contrast, the intra-transect comparison in the patchy OD meadow shows little evidence of attenuation in TOC or BC across the meadow. Nevertheless, there is evidence of resuspension, as implied by the increased particle size and decreased silt-clay fraction away from the exposed edge. Additionally, particle size increased within patches over adjacent unvegetated areas. This pattern suggests that a sparse, patchy configuration allows sufficient space for the seagrass leaves to add to turbulence and resuspension (17). For the urban patchy OD meadow, BC’s relatively invariant response with TOC, and its high content is consistent with; 1) atmospheric deposition at the meadow scale or greater (8); and 2) the suggested greater levels of turbulence and resuspension that can result in a greater loss of litter and oxidation of the more labile fractions. Although, the loss of the labile fraction may have been tempered by a high edaphic TOC contents, the result of a more eutrophic polluted environ. Taken together, the atmospheric addition of relatively pure BC to the urban meadow over BC supplied and diluted by soil organics to the rural meadows is a likely reason for the consistently greater BC/TOC fractions. A difference that was likely compounded by the patchy state of the urban system over that of the more contiguous rural meadows.

## Conclusion

The study demonstrated that allochthonous recalcitrant BC may represent a major fraction of the sediment TOC content of coastal seagrass meadows, and its underestimation may lead to significant overestimates of the carbon sink services of such meadows. Within rural contiguous meadows, the size of seagrass canopy species appears to explain a higher than predicted BC contribution to sedimentary TOC. In contrast, in the urban meadow under land development pressure, BC content is largely independent of canopy parameters and TOC variability, and this meadow supports the highest and most consistent BC/TOC fractions. Clearly, more work is needed within Southeast Asian BC hotspots, both rural and urban, to understand the full extent of BC stock bias. We thus recommend moving away from simple mass balance estimation approaches towards one that includes stability and origin via incorporation of the concept of allochthonous recalcitrance, whether of BC or other recognised types.

Supporting data and figures and methods can be found in the electronic supplementary material. All the authors assisted in fieldwork and analysis of the samples. J.B.G. led the writing. C.C.H. compiled the supplemental material and the statistical analyses set within the tables. All authors approved the final version of the manuscript and agree to be accountable for all aspects of the manuscript. We declare we have no competing interests. Funding was provided by Sabah Park’s Tun Mustapha Park Scientific Expedition (SDK0006-2017), and Ministry of Science Technology and Innovation (FRGS0424-SG-1/2015).

## References

1. McLeod E, Chmura GL, Bouillon S, Salm R, Björk M, Duarte CM, et al. A blueprint for blue carbon: Toward an improved understanding of the role of vegetated coastal habitats in sequestering CO2. Frontiers in Ecology and the Environment. 2011;9(10):552–60.

2. Hill R, Bellgrove A, Macreadie PI, Petrou K, Beardall J, Steven A, et al. Can macroalgae contribute to blue carbon? An Australian perspective. Limnology and Oceanography. 2015;60(5):1689–706.

3. Duarte CM, Middelburg JJ, Caraco N. Major role of marine vegetation on the oceanic carbon cycle. Biogeosciences. 2005;2(1):1–8.

4. Christianen MJA, van Belzen J, Herman PMJ, van Katwijk MM, Lamers LPM, van Leent PJM, et al. Low-Canopy Seagrass Beds Still Provide Important Coastal Protection Services. PLoS ONE. 2013;8(5):e62413.

5. Kristensen E, Ahmed SI, Devol AH. Aerobic and anaerobic decomposition of organic matter in marine sediment: Which is fastest? Limnology and Oceanography. 1995;40(8):1430– 7

6. Kennedy H, Beggins J, Duarte CM, Fourqurean JW, Holmer M, Marba N, et al. Seagrass sediments as a global carbon sink: Isotopic constraints. Global Biogeochemical Cycles. 2010;24, GB4026.

7. Cathalot C, Rabouille C, Tisnérat-Laborde N, Toussaint F, Kerhervé P, Buscail R, et al. The fate of river organic carbon in coastal areas: A study in the RhÔne River delta using multiple isotopic (δ13C, δ14C) and organic tracers. Geochimica Et Cosmochimica Acta. 2013;118:33–55.

8. Chew ST, Gallagher JB. Accounting for black carbon lowers estimates of blue carbon storage services. Scientific reports. 2018;8(1):2553.

9. Gaveau DLA, Salim MA, Hergoualc’h K, Locatelli B, Sloan S, Wooster M, et al. Major atmospheric emissions from peat fires in Southeast Asia during non-drought years: evidence from the 2013 Sumatran fires. Scientific reports. 2014;4:6112.

10. Chiu S-H, Huang Y-H, Lin H-J. Carbon budget of leaves of the tropical intertidal seagrass Thalassia hemprichii. Estuarine, Coastal and Shelf Science. 2013;125:27–35.

11. Ricart AM, Dalmau A, Pérez M, Romero J. Effects of landscape configuration on the exchange of materials in seagrass ecosystems. Marine Ecology Progress Series. 2015;532:89–100.

12. Portillo E. Relation between the type of wave exposure and seagrass losses (Cymodocea nodosa) in the south of Gran Canaria (Canary Islands - Spain). Oceanological and Hydrobiological Studies. 2014;43(1):29–40.

13. Oksanen L. Logic of experiments in ecology: is pseudoreplication a pseudoissue? Oikos. 2001;94(1):27–38.

14. Kilminster K, McMahon K, Waycott M, Kendrick G, Scanes P, McKenzie L, et al. Unravelling complexity in seagrass systems for management: Australia as a microcosm. Science of The Total Environment. 2015;534:97–109.

15. Freeman AS, Short FT, Isnain I, Razak FA, Coles RG. Seagrass on the edge: Land-use practices threaten coastal seagrass communities in Sabah, Malaysia. Biological Conservation. 2008;141(12):2993–3005.

16. Fonseca MS, Cahalan JA. A preliminary evaluation of wave attenuation by four species of seagrass. Estuarine, Coastal and Shelf Science. 1992;35(6):565–76.

17. van Katwijk MM, Bos AR, Hermus DCR, Suykerbuyk W. Sediment modification by seagrass beds: Muddification and sandification induced by plant cover and environmental conditions. Estuarine, Coastal and Shelf Science. 2010;89(2):175–81.

