## Supplemental Figure S1, Tables S2, S3, S4 for "Carbon stocks of coastal seagrass in Southeast Asia may be far lower than anticipated when accounting for black carbon"

**Carbon stock services for tropical coastal seagrass can be far lower than anticipated when accounting for black carbon**

John B. Gallagher^1,2^, Chuan Chee Hoe^4^ Yap Tzuen-Kiat^3^, and Farahain W^3^

^1^Centre for Marine and Coastal studies, Universiti Sains Malaysia, 11800, Penang, Malaysia

^2^Institute for Marine and Antarctic Studies, University of Tasmania, 7000, Tasmania, Australia

^3^Borneo Marine Research Institute, Universiti Malaysia Sabah, 88400, Sabah, Malaysia

^4^Faculty of Science and Natural Resources, Universiti Malaysia Sabah, 88400, Sabah, Malaysia

Correspondence and requests for materials should be addressed to John B Gallagher

**Collection and processing of sediments**

A range of sedimentological and seagrass biological variables were measured within each quadrat (table 1) along with the first 1 cm of sediment for BC and TOC contents, taken in situ using a surface scoop. For each of the 2 transects at each location, 11 samples were taken, labelled 1-11 for the first transect and 12-22 for the second transect. Samples of bare sediments were also taken for BC and TOC contents. Sampling across the meadows was performed within quadrats laid down every 5 m along two parallel 50 m transects, and perpendicular to the exposed boundary of a prevailing Monsoon (Supplementary Figure S1). The intra-transect sites being spaced to capture the patch variability across that portion of the meadow, and the distance between transects to determine difference in the central statistic at the meadow scale (50 m) (15). The sites were selected away from adjacent shallow banks of coral rubble, and windward to the prevailing Monsoon, along with an additional 10 samples along the central channel adjacent to seagrass meadows of Salut–Mengkabong estuary. For details of sample treatment, sediment particle size distribution and analysis of TOC and BC as measured using a gravimetric chemo-thermal oxidation protocol can be found elsewhere (5).

**Data analysis of seagrass meadow variation in between transects.**

All statistical analysis, t-tests of difference between means of variables and parameters were calculated in SigmaPlot. Where normality was violated, a Mann-Whitney test for differences between medians was used. Where differences in the means were visually apparent but variability gave immediately outside P > 0.05, a power analysis (80 %) was used to indicate how many additional samples would be require to resolve a possible Type 1 error (SigmaPlot). Similarities between least squares linear regression parameters were tested using ANCOVA within PAST.

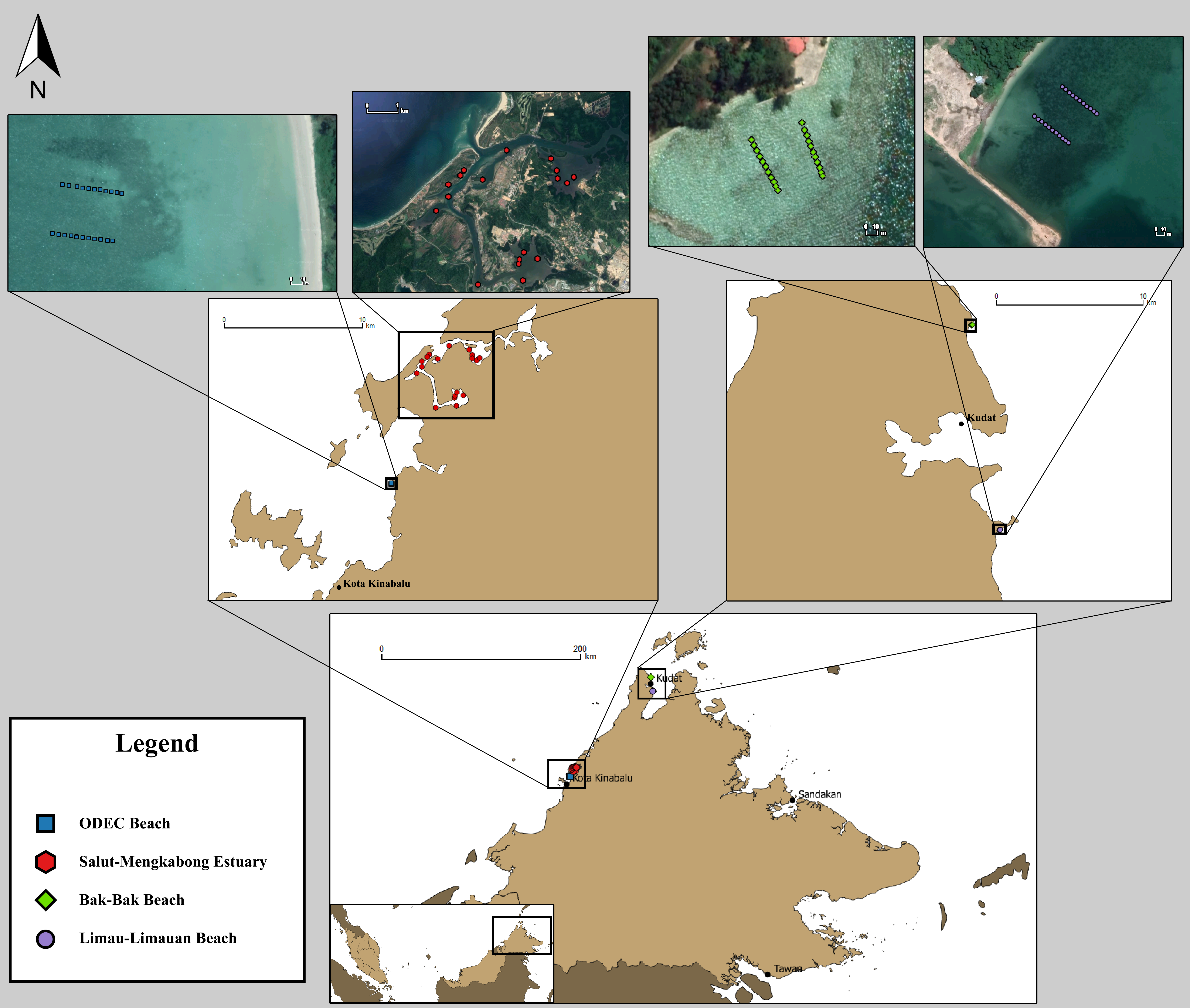

**Supplemental Figure 1:** Map of Sabah, showing the locations of all transects taken during the study and their position in the state. Top: Satellite images of study quadrats, with sampling sites labelled respectively. Left to right: ODEC Beach, Salut-Mengkabong Estuary, Bak-Bak Beach and Limau-Limauan beach. Map data: Google, 2018; Landsat/Copernicus, Digital Globe, Data SIO, NOAA, U.S. Navy, NGA, GEBCO, TerraMetrics and Bornean Biodiversity & Ecosystems Conservation (BBEC) Sabah and WWF Malaysia, 2017. The linemap was produced with with QGIS v2.18.14, Adobe Illustrator CS6 and Adobe Photoshop CS6.

**Supplementary Table S1.** ODEC Beach seagrass sediment sample positions, Total Organic Carbon and Black Carbon percentages, percentage of BC to TOC content of the sediments sampled at the study sites, along with the seagrass quadrants’ sedimentological and seagrass biological variables. Sites labelled with the prefix T1 belong to the inner transect, while sites labelled with T2 belong to the outer transect.

| **Site ID** | **Coordinates** | **TOC**  **(% dry mass)** | **BC**  **(% dry mass)** | **BC/TOC**  **(%)** | **Seagrass Coverage (%)** | **Canopy Height (cm)** | **Particle Size (µm)** | **Silt-clay Fraction (%)** |
| --- | --- | --- | --- | --- | --- | --- | --- | --- |
| T1-OD1 | 6.048259 N  116.109457 E | 0.80 | 0.24 | 30.16 | 13 | 19.5 | 66.77 | 18.83 |
| T1-OD2 | 6.048263 N  116.109415 E | 0.96 | 0.31 | 32.16 | 13 | 17.7 | 105.82 | 7.93 |
| T1-OD3 | 6.048269 N  116.109366 E | 1.24 | 0.30 | 24.38 | 38 | 20.1 | 94.64 | 14.51 |
| T1-OD4 | 6.048273 N  116.109322 E | 0.99 | 0.27 | 27.54 | 0 | 0.0 | 99.69 | 10.12 |
| T1-OD5 | 6.048279 N  116.109275 E | 1.32 | 0.31 | 23.70 | 38 | 9.8 | 99.31 | 11.54 |
| T1-OD6 | 6.048285 N  116.109230 E | 1.02 | 0.29 | 28.12 | 13 | 2.6 | 94.83 | 11.61 |
| T1-OD7 | 6.048291 N  116.109184 E | 0.96 | 0.27 | 28.68 | 13 | 16.4 | 94.07 | 14.50 |
| T1-OD8 | 6.048298 N  116.109142 E | 1.28 | 0.34 | 26.33 | 0 | 0.0 | 103.69 | 11.44 |
| T1-OD9 | 6.048302 N  116.109096 E | 1.33 | 0.29 | 21.75 | 0 | 0.0 | 98.19 | 12.88 |
| T1-OD10 | 6.048307 N  116.109050 E | 0.92 | 0.26 | 28.08 | 0 | 0.0 | 92.09 | 14.75 |
| T1-OD11 | 6.048314 N  116.109004 E | 1.04 | 0.31 | 29.76 | 0 | 0.0 | 89.45 | 13.44 |
| T2-OD12 | 6.048617 N  116.109525 E | 0.90 | 0.25 | 28.06 | 0 | 0.0 | 121.72 | 3.34 |
| T2-OD13 | 6.048624 N  116.109486 E | 0.96 | 0.29 | 29.68 | 0 | 0.0 | 114.76 | 4.73 |
| T2-OD14 | 6.048628 N  116.109443 E | 0.94 | 0.32 | 34.11 | 0 | 0.0 | 115.70 | 6.17 |
| T2-OD15 | 6.048634 N  116.109404 E | 1.04 | 0.29 | 27.46 | 0 | 0.0 | 118.60 | 5.61 |
| T2-OD16 | 6.048643 N  116.109361 E | 1.17 | 0.29 | 24.56 | 0 | 0.0 | 128.11 | 5.49 |
| T2-OD17 | 6.048647 N  116.109319 E | 1.10 | 0.30 | 27.49 | 0 | 0.0 | 123.11 | 6.40 |
| T2-OD18 | 6.048651 N  116.109275 E | 1.00 | 0.31 | 31.11 | 13 | 14.5 | 136.26 | 4.13 |
| T2-OD19 | 6.048657 N  116.109230 E | 0.96 | 0.32 | 33.07 | 0 | 0.0 | 127.11 | 5.85 |
| T2-OD20 | 6.048665 N  116.109189 E | 0.95 | 0.29 | 30.38 | 0 | 0.0 | 125.72 | 4.58 |
| T2-OD21 | 6.048675 N  116.109128 E | 1.06 | 0.31 | 29.19 | 38 | 9.4 | 140.10 | 4.37 |
| T2-OD22 | 6.048680 N  116.109077 E | 1.50 | 0.29 | 19.51 | 38 | 17.2 | 135.12 | 5.24 |

**Supplementary Table S2.** Bak-Bak Beach seagrass sediment sample positions, Total Organic Carbon and Black Carbon percentages, percentage of BC to TOC content of the sediments sampled at the study sites, along with the seagrass quadrants’ sedimentological and seagrass biological variables. Sites labelled with the prefix T1 belong to the inner transect, while sites labelled with T2 belong to the outer transect.

| **Site ID** | **Coordinates** | **TOC**  **(% dry mass)** | **BC**  **(% dry mass)** | **BC/TOC**  **(%)** | **Seagrass Coverage (%)** | **Canopy Height (cm)** | **Particle Size (µm)** | **Silt-clay Fraction (%)** |
| --- | --- | --- | --- | --- | --- | --- | --- | --- |
| T1-BB1 | 6.944166 N 116.839410 E | 0.77 | 0.26 | 34.36 | 38 | 5.3 | 115.34 | 4.96 |
| T1-BB2 | 6.944124 N 116.839431 E | 0.78 | 0.35 | 44.68 | 38 | 6.0 | 124.12 | 3.90 |
| T1-BB3 | 6.944081 N 116.839454 E | 0.72 | 0.28 | 39.15 | 38 | 4.7 | 118.48 | 4.44 |
| T1-BB4 | 6.944034 N 116.839476 E | 0.56 | 0.22 | 39.28 | 63 | 5.3 | 118.81 | 4.19 |
| T1-BB5 | 6.943989 N 116.839501 E | 0.67 | 0.26 | 38.92 | 63 | 7.0 | 121.67 | 5.76 |
| T1-BB6 | 6.943952 N 116.839522 E | 0.90 | 0.34 | 37.97 | 63 | 6.7 | 141.80 | 2.86 |
| T1-BB7 | 6.943911 N 116.839544 E | 0.87 | 0.33 | 38.18 | 63 | 9.3 | 128.63 | 5.07 |
| T1-BB8 | 6.943866 N 116.839566 E | 0.86 | 0.32 | 37.72 | 0 | 0.0 | 138.73 | 3.12 |
| T1-BB9 | 6.943830 N 116.839589 E | 0.86 | 0.32 | 36.69 | 13 | 8.3 | 139.35 | 2.89 |
| T1-BB10 | 6.943796 N 116.839605 E | 0.83 | 0.30 | 36.33 | 0 | 0.0 | 133.41 | 3.34 |
| T1-BB11 | 6.943765 N 116.839620 E | 0.58 | 0.19 | 31.77 | 0 | 0.0 | 137.12 | 3.36 |
| T2-BB12 | 6.944300 N 116.839809 E | 0.48 | 0.17 | 35.90 | 63 | 2.0 | 130.59 | 4.59 |
| T2-BB13 | 6.944243 N 116.839832 E | 0.69 | 0.24 | 34.77 | 63 | 2.0 | 131.49 | 5.07 |
| T2-BB14 | 6.944201 N 116.839848 E | 0.54 | 0.16 | 30.21 | 63 | 2.2 | 141.69 | 3.93 |
| T2-BB15 | 6.944155 N 116.839864 E | 0.65 | 0.23 | 35.91 | 13 | 2.3 | 142.72 | 3.32 |
| T2-BB16 | 6.944114 N 116.839882 E | 0.74 | 0.25 | 33.51 | 63 | 2.0 | 138.63 | 4.45 |
| T2-BB17 | 6.944069 N 116.839901 E | 0.72 | 0.23 | 31.19 | 63 | 1.5 | 136.01 | 4.92 |
| T2-BB18 | 6.944028 N 116.839917 E | 0.73 | 0.27 | 36.73 | 0 | 0.0 | 143.08 | 3.52 |
| T2-BB19 | 6.943987 N 116.839932 E | 0.71 | 0.23 | 31.87 | 13 | 4.3 | 140.24 | 2.90 |
| T2-BB20 | 6.943943 N 116.839949 E | 0.63 | 0.19 | 30.73 | 63 | 5.3 | 128.13 | 5.01 |
| T2-BB21 | 6.943906 N 116.839963 E | 0.74 | 0.25 | 33.53 | 63 | 5.7 | 126.82 | 4.64 |
| T2-BB22 | 6.943877 N 116.839975 E | 0.65 | 0.23 | 35.06 | 88 | 4.7 | 133.72 | 3.70 |

**Supplementary Table S3.** Limau-Limauan Beach seagrass sediment sample positions, Total Organic Carbon and Black Carbon percentages, percentage of BC to TOC content of the sediments sampled at the study sites, along with the quadrants’ sedimentological and seagrass biological variables. Sites labelled with the prefix T1 belong to the inner transect, while sites labelled with T2 belong to the outer transect.

| **Site ID** | **Coordinates** | **TOC**  **(% dry mass)** | **BC**  **(% dry mass)** | **BC/TOC**  **(%)** | **Seagrass Coverage (%)** | **Canopy Height (cm)** | **Particle Size (µm)** | **Silt-clay Fraction (%)** |
| --- | --- | --- | --- | --- | --- | --- | --- | --- |
| T1-LL1 | 6.818167 N 116.856433 E | 0.77 | 0.33 | 42.50 | 0 | 0.0 | 21.22 | 75.23 |
| T1-LL2 | 6.818134 N 116.856472 E | 0.26 | 0.08 | 33.10 | 38 | 18.0 | 77.71 | 13.40 |
| T1-LL3 | 6.818104 N 116.856510 E | 0.25 | 0.07 | 26.48 | 13 | 10.3 | 73.35 | 14.92 |
| T1-LL4 | 6.818075 N 116.856548 E | 0.31 | 0.06 | 19.13 | 13 | 8.0 | 75.69 | 14.37 |
| T1-LL5 | 6.818047 N 116.856585 E | 0.15 | 0.03 | 22.56 | 0 | 0.0 | 66.69 | 15.41 |
| T1-LL6 | 6.818017 N 116.856620 E | 0.24 | 0.05 | 21.17 | 63 | 18.7 | 63.74 | 16.92 |
| T1-LL7 | 6.817989 N 116.856657 E | 0.26 | 0.08 | 29.06 | 38 | 5.5 | 94.40 | 8.29 |
| T1-LL8 | 6.817958 N 116.856694 E | 0.28 | 0.06 | 22.02 | 63 | 18.7 | 84.30 | 11.51 |
| T1-LL9 | 6.817931 N 116.856729 E | 0.38 | 0.11 | 29.86 | 63 | 10.0 | 52.67 | 21.52 |
| T1-LL10 | 6.817908 N 116.856759 E | 0.32 | 0.15 | 48.32 | 63 | 9.8 | 69.71 | 15.83 |
| T1-LL11 | 6.817886 N 116.856791 E | 0.77 | 0.20 | 25.92 | 63 | 9.0 | 72.72 | 13.93 |
| T2-LL12 | 6.818475 N 116.856724 E | 0.14 | 0.05 | 32.42 | 13 | 8.7 | 81.93 | 12.13 |
| T2-LL13 | 6.818446 N 116.856761 E | 0.23 | 0.03 | 12.96 | 38 | 18.0 | 66.07 | 17.27 |
| T2-LL14 | 6.818414 N 116.856799 E | 0.24 | 0.05 | 21.46 | 38 | 14.3 | 74.48 | 14.77 |
| T2-LL15 | 6.818384 N 116.856836 E | 0.16 | 0.03 | 21.28 | 38 | 16.7 | 47.79 | 24.22 |
| T2-LL16 | 6.818357 N 116.856873 E | 0.15 | 0.00 | 2.13 | 38 | 21.0 | 76.62 | 14.75 |
| T2-LL17 | 6.818329 N 116.856910 E | 0.16 | 0.01 | 4.93 | 38 | 17.3 | 60.68 | 19.52 |
| T2-LL18 | 6.818300 N 116.856949 E | 0.32 | 0.10 | 30.81 | 63 | 19.7 | 56.46 | 22.06 |
| T2-LL19 | 6.818270 N 116.856986 E | 0.28 | 0.08 | 28.67 | 13 | 9.0 | 96.68 | 9.71 |
| T2-LL20 | 6.818245 N 116.857024 E | 0.30 | 0.07 | 23.02 | 13 | 5.0 | 102.30 | 8.11 |
| T2-LL21 | 6.818217 N 116.857057 E | 0.11 | 0.05 | 42.20 | 13 | 6.7 | 113.58 | 6.82 |
| T2-LL22 | 6.818192 N 116.857091 E | 0.44 | 0.15 | 34.84 | 13 | 9.0 | 117.19 | 6.34 |
